# Interplay between the Xer recombination system and the dissemination of antibioresistance in *Acinetobacter baumannii*

**DOI:** 10.1101/2024.04.09.588662

**Authors:** Blanchais Corentin, Pages Carine, Manuel Campos, Kenza Boubekeur, Contarin Rachel, Mathias Orlando, Siguier Patricia, Laaberki Maria-Halima, Cornet François, Charpentier Xavier, Rousseau Philippe

## Abstract

Antibiotic-resistant infections pose a pressing challenge in clinical settings. Plasmids are widely recognized for hastening the emergence of resistance by facilitating horizontal gene transfer of antibiotic resistance genes among bacteria. We explore this inquiry in *Acinetobacter baumannii*, a globally emerging nosocomial pathogen responsible for a wide array of infections with worrying accumulation of resistances, notably involving plasmids. In this specie, plasmids of the Rep_3 family harbor adaptive genes within variable regions edged by potential site-specific recombination sites recognized by the XerCD recombinase. We first show that the Xer system of *Acinetobacter baumannii* functions as described in *Escherichia coli*, resolving chromosome dimers at the *dif* site as well as recombining plasmid-borne sites. The multiple Xer recombination sites found in Rep_3 plasmids do not, however, allow excising plasmid fragments. They rather recombine to co-integrate plasmids, which may then further evolve to exchange genes. Co-integrates represent a significative part of the plasmid population and their formation is controlled by the sequence of the recombination sites determining their compatibility between the recombining sites. We conclude that plasmids frequently exchange genes in *Acinetobacter baumannii* using Xer recombination, allowing a high level yet controlled plasticity involved in the acquisition and combination of resistance genes.

## Introduction

In bacteria, most replicons are circular and during their replication, an odd number of homologous recombination events leads to their dimerization. This impairs the even distribution of replicons between daughter cells (segregation) during cell division. To resolve this problem, prokaryotes have evolved replicon dimer resolution sites acted on by dedicated Xer recombinases (XerC and XerD in most cases)(1, 2), that we call *xrs* (Xer Recombination sites). The importance of this function for faithful replicon segregation explains its high conservation and the Xer system is now considered as one of the most conserved structural features of circular chromosomes in Bacteria and Archaea (3, 4).

The XerCD recombination process is particularly well described for the unique *xrs* located on bacterial chromosomes, known as the *dif* site (Figure 1A). The *dif* site is composed of two protein-binding arms separated by a central region. XerC and XerD respectively bind specifically to their binding arm (5) and two XerCD-*dif* complexes then interact to form the XerCD-*dif* synapse. Within this synapse, only one type of recombinase, either XerC or XerD, is expected to be active and each of the two units of that recombinase cuts the DNA strand at the *dif* site to which it is bound (5). This nucleophilic attack of DNA, mediated by a conserved tyrosine residue, forms a covalent link between the recombinase and the *dif* site. The second step of the reaction is a strand exchange between the two *dif* copies in the central region followed by ligation, which creates a Holliday junction. This intermediate isomerizes, thereby activates the second pair of recombinases, which cut and exchange the second pair of strands, finishing the recombination reaction (5). In this process, the two pairs of recombinases are sequentially activated to catalyze the exchange of the two DNA strands (6). Therefore, the selection of the first active pair of recombinases controls the reaction and it has been proposed that, within the XerCD-*dif* synapse, XerC is the one initially active whilst XerD is initially inactive. Consequently, the reaction is blocked at the Holliday Junction step and thus tends to be reversible, without recombination (7). To catalyze a complete recombination process, XerD must be activated by FtsK, a division septum-associated DNA translocase, which is essential for cell division (8–10). By recognizing KOPS sequences, which are oriented toward *dif* on each chromosome replichores, FtsK translocates toward *dif* (11–13). Upon reaching the *dif* site, FtsK activates XerCD-*dif* recombination through a specific contact with the carboxy-terminal part of XerD (9). This FtsK control of the XerCD-*dif* recombination permits a spatio-temporal control of chromosome dimer resolution: at midcell during septation.

**Figure 1:**
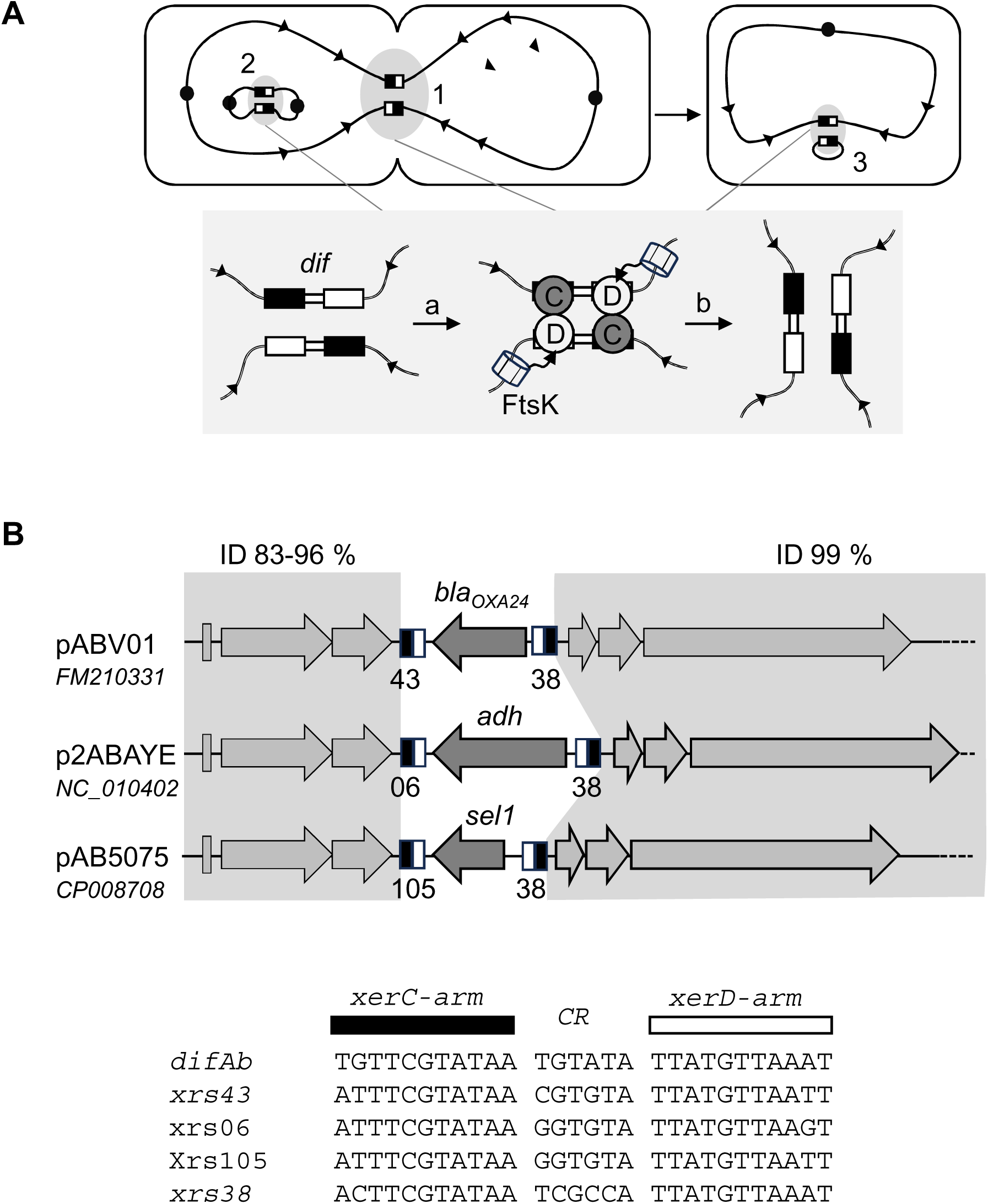
Xer System. **A)** Xer is a site-specific recombination system that resolves chromosome dimers before cell division (1). Xer is hijacked by plasmids to resolves their dimers, ensuring their correct vertical transmission (2) and by IMEX to ensure their integration to the chromosome (3). Xer recombination sites (*xrs*) are represented as black&whight boxes. The *xrs* present on the chromosome is called *dif.* Black circles represent replication origins. The *xrs* are composed of a XerC binding arm (black rectangle) and a XerD binding arm (wight rectangle) separated by a central region (CR). XerC (grey circle) and XerD (wight circle) binds *dif* (or *xrs*) and assemble a synaptic complex (a) within which, once activated by FtsK, they will catalyze the successives exchanges of the two DNA strands of the central region (b) to form the recombined DNA. **B)** Some *Ab* plasmids carry multiple *xrs* (B&W boxes) and are highly identical (ID) except in regions flanked by *xrs*

Plasmids are also affected by this dimerization problem (14). Most of them use the Xer system to resolve their dimeric forms and some have evolved *xrs* to be independent of the bacterial cell cycle (e.g.: *cer* or *psi* are FtsK-independent *xrs*; for review (15)).

While it is expected that plasmid carry a single *xrs* site to resolve dimers (15), an increasing number of plasmids harboring multiple *xrs* has been described (called *Re27*, *pdif* or *pXerC/D*, (16–20)). The functional significance of xrs multiplicy remains unresolved. Most cases are found in *Acinetobacter baumannii* (*Ab*), a human pathogen involved in nosocomial diseases which are difficult to treat due their resistance to multiple antibiotics. For instance, the pABV01 plasmid contains two *xrs* flanking the *bla_oxa-24_* gene that confers resistance to carbapenems (17). This plasmid is related to two others (p2ABAYE and pAB0057) that are very similar but for the region contained between the two *xrs* (Fig. 1B). From these observations, emerged the concept of *xrs-* cassettes (adaptative genes flanked by two *xrs*) suggesting the implication of Xer system in antibiotic resistant genes (ARG) dissemination (also called *pdif*-modules; (18, 20, 21). Recent work clearly suggests that *xrs* present on *Ab* plasmids are hot spot of recombination between plasmids (19, 22, 23) and that they may be processed by Xer recombinases (24). However, very little is known about the activity of these sequences and/or cassettes and whether they are *bona fide* xer recombination sequences. We present here a functional study of the Xer system of *Ab* on chromosomal (*dif*) and plasmidic *xrs.* This allowed us to test models of mobility of *xrs*-cassettes. Our results support a model of gene exchange between *Ab* plasmids via an original cointegration/resolution mechanism.

## Material and Methods

### *Escherichia coli* strains and plasmids

Cassette carrying plasmids were derived from pFX346 (25). Tandem of *xrs* (direct or inverse orientation depending on the cases) separated by a SalI site were first obtained by hybridization of two complementary oligonucleotides and inserted into pFX346 (between BamHI and SphI). *lacIq* was then inserted between *xrs* (SalI) to give pROUT29 (*dif_Ab_*-*lacIq*-*dif_Ab_*), pROUT25 (*xrs43*-*lacIq*-*xrs38_inv_*), pROUT31 (*xrs105*-*lacIq*-*xrs38_inv_*) and pROUT32 (*xrs6*-*lacIq*-*xrs38_inv_*). *E.coli* strains were derived from K12 strain LN2666. CP1518 strain was derived from CP109 (LN2666 Δ*dif*::Tc, Δ(*lacI*), *xer*::Gm)(26), by insertion of a *dif_Ab_-lacI-dif_Ab_* cassette into the Tc gene (insertion-excision, Cornet *et al.*, 1996). To do this modification, the *dif_Ab_-lacI-dif_Ab_* cassette was first inserted between BamH1 and SpaI sites within plasmid pLN135 to give pCP179. The *recF*::Kn allele (KEIO collection (27)) was introduced (P1 transduction) in CP1088 (LN2666 *ΔxerD*, *ΔxerC* (28)) to give CB50 (LN2666 *ΔxerD*, *ΔxerC*, *recF*::Kn). *A. baumannii xer* genes (*xerC_Ec_, xerC_Ab_, xerD_Ec_, xerD_Ab_, xerC_RQ_, xerD_RQ_* and *xerDγ_ftsK_*) were ordered from GenScript compagnie. There where cloned into pET32 to give production plasmids pROUT26 (*xerC_Ab_*), pROUT27 (*xerD_Ab_*), pCB01 (*xerC_Ab_*-R254Q), pCB02 (*xerD_Ab_-R255Q*) and pCB03 (*xerDγ_ftsK_*). These genes were also used to constructed pSC101 based in vivo expression vectors. The pBAD promotor (pBAD18) was cloned into pSC101 plasmid (29) and *xer* genes where cloned alone, or in operon (In-Fusion cloning), under the control of pBAD promoter to give pLN1xerCD_Ec_, pLN1*xerCD_Ab_*, pLN1*xerCDγ_Ab_* and pLN1*xerCD_RQ_*.

### *Acinetobacter baumannii* strains and plasmids

Ab5075-T and AYE-T are *Acinetobacter baumannii* stains derived from AB5075-UW (NZ_008706) and AYE (NC_010410) (30). To obtain CB-AB01 (Ab5075-T *xerC*::*sacB_aacC*) and CB-AB03 (ABAYE-T *xerC*::*sacB_aacC4*), AYE-T and AB5075-T were transformed with DNA fragments carrying a *sacB_aacC4* gene flanked by two 2kb homology regions. This DNA fragment was obtained by cross-over-PCR assembly of 3 DNA fragments. Oligonucleotides used are 74-75, 78-79 and 76-77 (TableSup1). Transformants were plated on Luria Broth (LB) agar supplemented with 30 μg/mL apramycin. These strains were then transformed with a DNA fragment containing the two 2kb homology (Oligonucleotides 74-80 and 78-81, TableSup1) to obtained ABAYEΔ*xerC* and AB5075Δ*xerC*. Transformants were selected on M9 agar complemented with 10% sucrose (30). Plasmids used in this work are pABV01(FM210331), p1ABAYE (NC_010401) p2ABAYE(NC_010402) and p2AB5075(CP008708).

### In vitro experiments

Xer proteins were purified as described previously (25, 28). To do EMSA experiments, 28-bp 5′-end-labeled [CY3] DNA fragments carrying *dif_Ab_* or *xrs* sites were obtained by hybridization of complementary strands (TableSup1). EMSA reactions were performed as previously described (25, 28, 31) and analyzed with a typhoon TRIO GE. Cleavage assay were performed as previously described (32)(see Table Sup). DNA substrates used were produced by hybridization of different oligonucleotides (TableSup1) and results of the assay were analyzed on SDSPAGE (Mini-PROTEAN TGX gels – 4-20%).

### *In vivo* recombination assay

For chromosome-based deletion assay, CP1518 was transformed with pLN1-type expression vector (pLN1xerCD_Ec_, pLN1xerCD_Ab_, pLN1xerCDγ_Ab_ and pLN1xerCD_RQ_). Serial dilution of the transformants were plate on LB agar medium complemented with 5µg/mL chloramphenicol, 40µg/mL XGal and 0.4% Arabinose or Glucose and grown overnight at 37°C as previously describe (31).

For plasmid-based deletion assay, CB50 was transformed with pROUT29, pROUT25, pROUT31 or pROUT32, together with a pLN1-based xer expression plasmid. Transformants were plated on LB agar complemented with 5µg/mL chloramphenicol and 25µg/mL ampicillin. After overnight culture in selective LB medium with 0.4% arabinose, plasmid content was extracted (QIAprep Spin Miniprep Kit) and analyzed on 1% agarose gel electrophoresis.

### Cointegrates detection and quantification

Cointegrates were detected by PCR (EmeraldAmp GT PCR Master Mix) on DNA plasmid matrices purified from *A. baumannii* strains (QIAprep Spin Miniprep Kit).

Pairs of oligonucleotides used to detect non-recombined plasmids were: 94-105 for *xrs105*; 95-101 for *xrs38* and 103-121 for *xrs10*. The expected lengths of the amplicons were 1.3kpb, 1.0kpb and 1.2kpb respectively. Pairs of oligonucleotides used to detect cointegrates were: 94-117 for *xrs6/105* recombination; 95-119 for *xrs66/38* recombination and 103+120 for *xrs107/10* recombination. The expected lengths of the amplicons were 0.8kpb, 0.7kpb and 1.2kpb respectively. PCR products were analyzed on 1% agarose gel electrophoresis.

Cointegrates quantification was performed using droplet digital PCR (Bio-Rad QX200 droplet-reader). Oligonucleotides were designed by using Primer3Plus program (Figure SUP1). Matrix DNA was extracted from 1mL of OD_600_ = 0.6 cultures (Wizard Genomic DNA Purification Kit). DNA was then digested with EcoRI (FastDigest ThermoFisher) and diluted 10^2^ time. A typical 20μl ddPCR reaction contains 1µL of prepared gDNA, 2X EvaGreen ddPCR Supermix (Bio-Rad) and 50nM of each primer. A typical amplification reaction is: 5 min at 95°C; 40x (30 sec. at 95°C ; 30 sec. at 58°C; 1 min. at 72°C); storage at 4°C. A 2.5°C/sec. ramp rate was used. Results were analysed using QuantaSoft droplet reader software.

## Results

### Characterization of Xer system of *A. baumannii*

We first investigate the functionality of *A. baumannii*’s Xer system (Xer_Ab_). To this end, full-length XerC and XerD proteins from strain AB5075 (subsequently referred to as XerC_Ab_ and XerD_Ab_) were purified in heterologous system. Binding of purified proteins to their expected substrate, *dif_Ab_*-containing DNA, was assessed using Electrophoretic Mobility Shift Assay (EMSA, Fig. 2). Increasing concentration of XerC_Ab_ led to two shifted bands (C1 and C2). Based on previous observations in *E. coli* Xer system (Blakely), we inferred that these bands correspond to XerC_Ab_ bound to its binding site (C1) and to both XerC_Ab_ and XerD_Ab_ binding sites (C2). Equivalent results were obtained for XerD_Ab_. XerD_Ab_ appeared to bind *dif_Ab_* with higher affinity than XerC_Ab_. The C1 complexes migrates differently depending on the protein used, suggesting different conformation of these complexes (*e.g.* DNA bending (25)). When both XerC_Ab_ and XerD_Ab_ were added, only a complex equivalent to C2 was observed. We inferred that this complex corresponds to the binding of both XerC_Ab_ and XerD_Ab_ on their binding sites on *dif_Ab_*. Taken together, these results show that that Xer recombinases of *Ab* form a tripartite complex with *dif_Ab_*, as described for the *E. coli* system (5, 28).

**Figure 2:**
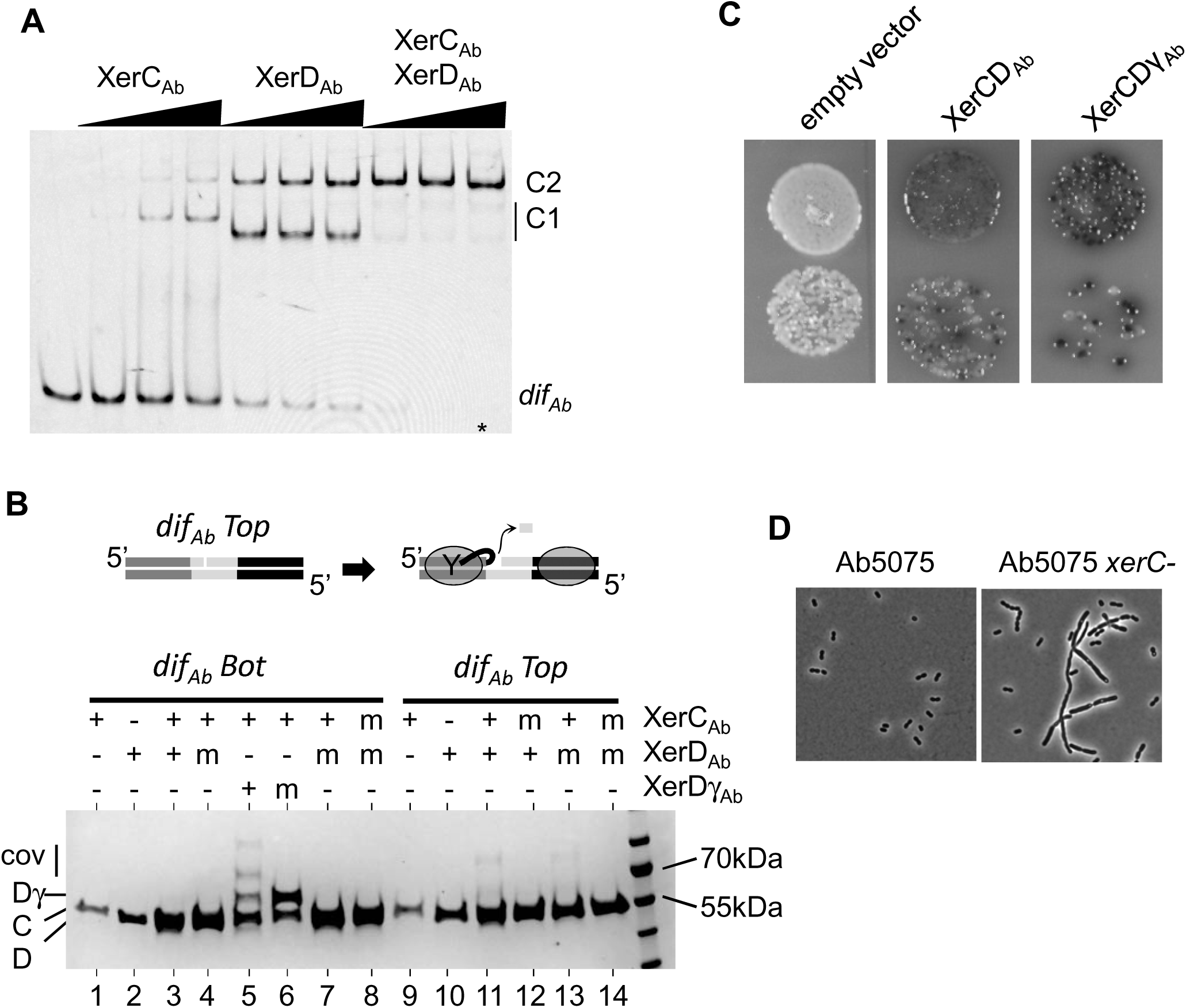
Activity of the Xer Systrem of *Ab*. **A)** EMSA of *dif_Ab_* by XerCD_Ab_. The DNA fragment (*dif_Ab_*) is 56 bp long and is 5’-CY3. XerC_Ab_ (mM) = 0; 0.48; 1.2 and 2.4. XerD_Ab_ (mM) = 0; 0.64; 1.6 and 3.2 **B)** SDS-PAGE analysis of suicide substrate assays (top diagram). DNA molecules (*dif_Ab_* Top or Bottom nicked) are 10mM whereas proteins are 2 mM. XerC_Ab_ is indicated as C, XerD_Ab_ as D, XerDg_Ab_ as Dg and covalent products made between DNA and proteins as cov **C).** *E. coli* (CP1518, *Δdif::difAb-lacI-difAb,Δ(lacI), xer::Gm*) was transformed with vector expressing *xerCDAb*, or *xerCD*γ_Ab_. Serial dilution of the transformant were plate on solid media complemented with XGal and Arabinose (M&M). Blue colonies correspond to Xer induced deletion of the lacI gene **D)** Micrographs of bacteria carrying, or not, a deletion of *xerC*. Strains were observed under phase-contrast microscopy (M&M).

We then checked if XerC_Ab_ and XerD_Ab_ can catalyze the first step of DNA recombination on *dif_Ab_*, i.e., DNA cleavage by the catalytic tyrosine residue, leading to the formation of a covalent link between the cleaved DNA and the Xer protein. To do so, we used a “suicide assay” into which the covalent DNA-protein intermediate is trapped (32)(Materials & Methods; Figure 2B). We observed that XerC_Ab_ cleaved *dif_Ab_*-containing DNA when in presence of XerD_Ab_ (Fig. 2B; lanes 9, 10 & 11). This cleavage was not observed when a catalytic residue of XerC_Ab_ was mutated (lane12). In contrast, XerD_Ab_ did not cleave *dif_Ab_*, even if in presence of XerC_Ab_ (Fig. 2B; lanes 1, 2 & 3). This later result suggests that, as in *E. coli*, activation by FtsK is required for cleavage by XerD_Ab_ (9). As expected, C-terminal fusion of AB5075 FtsKγ-domain to XerD, referred as XerDγ_Ab_, enables cleavage of *dif_Ab_*-containing DNA (Fig. 2B; lane 5).

To further characterize the Xer_Ab_ system, we replaced the *dif_Ec_* site on the *E. coli* chromosome by a *dif_Ab_-lacI-dif_Ab_* cassette in a strain deleted for the *xerC* and *lacI* genes (26)(Material & Methods). The resulting strain made white colonies when plated on LB in presence of the beta-galactosidase substrate (Xgal), LacI effectively repressing the beta-galactosidase-encoding gene *lacZ* (Fig. 2C, empty vector). When this strain was transformed with a plasmid expressing both *xerC_Ab_* and *xerD_Ab_* genes, blue colonies appeared, indicating that recombination events occurred (Fig. 2C, Table Sup2). Recombination was 5 time more efficient when *xerDγ_Ab_* was substituted to *xerD_Ab_* (Fig. 2C, Table Sup2), further demonstrating that Xer_Ab_ recombination is controlled by FtsK. To bring evidence of Xer*_Ab_* functionality in *A. baumannii* replication, we deleted the *xerC_Ab_* gene in strain AB5075. The resulting strain formed filament (Fig. 1D), similarly to observations performed in *E. coli* strains inactivated for the resolution of chromosome dimers do (33).

Taken together, all these results demonstrate that the XerCD/*dif* system of *A. baumannii* is canonical, behaving as site-specific recombination system involved in chromosome dimer resolution under the control of the division protein FtsK.

### Plasmids of *A. baumannii* carry multiple Xer Recombination Sites

We next checked if the *xrs* found in multiple copies in Ab plasmids are active. To do so we studied a set of *xrs* from three related plasmids of the Rep_3 family reported in clinical strains: pABV01; p2ABAYE; and p2AB5075. Results are summarized in Fig. 3A (Fig. Sup1 for raw data). All sequences assayed, but *xrs107* which is strongly degenerated, were cooperatively bound by XerC_Ab_ and XerD_Ab_. XerC_Ab_ alone bond poorly to these sequences whereas XerD_Ab_ exhibited a better apparent affinity. This is consistent with the fact that the predicted *xerC*-binding sites of these *xrs* is more divergent from the *dif* consensus than their *xerD-*binding sites (21). Most *xrs* were cleaved by XerC_Ab_ and XerDγ_Ab_ but not by XerD_Ab_ (Fig. 3, *xrs6* is presented as a typical example). We conclude that most *xrs* found in *Ab* plasmid appear active for Xer recombination.

**Figure 3:**
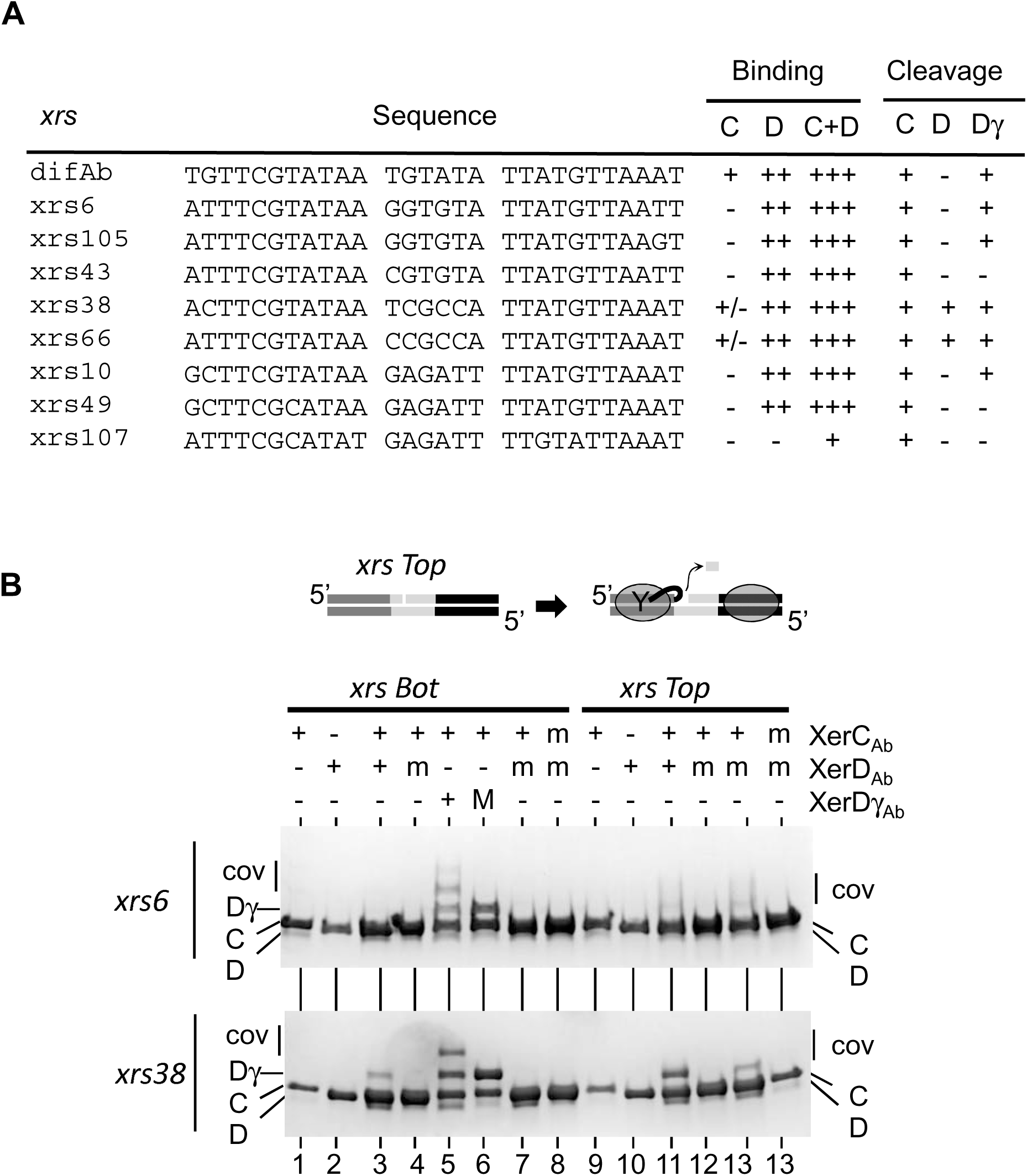
Activity of *xrs* sequences present on *Ab* plasmids. **A)** summary of EMSA and cleavage assays performed on different *xrs*. pABV01 contain *xrs38*, *xrs43* and *xrs10*. P1ABAYE contains *xrs*06, *xrs107* and *xrs66*. P2ABAYE contains *xrs38*, *xrs105* and *xrs49*. P2AB5075 contains *xrs38*, *xrs06* and *xrs49*. Experiment were performed like in figure 2. Raw data are presented in figures SUP1&2. **B)** SDS-PAGE analysis of cleavage assays (top diagram) on *xrs6* and *xrs38*. DNA molecules (*xrs* Top or Bottom nicked) are 10mM whereas proteins are 2 mM. XerC_Ab_ is indicated as C, XerD_Ab_ as D, XerDg_Ab_ as Dg and covalent products made between DNA and proteins as cov. Note that the covalent product XerD_Ab_-xrs38 migrate like XerDgAb (Lane 3 bottom gel).

Interestingly, *xrs38*, which is the only *xrs* to be present on the three plasmids presented on Fig. 1B, displays a different property. Indeed, we found that *xrs38* is bound and cleaved by XerC_Ab_ but also, and unexpectedly by XerD_Ab_ (Fig. 3B). Likewise, x*rs66*, closely related to *xrs38* but present on the plasmid p1ABAYE (see below) also exhibit this property. This suggests that *xrs38 and xrs66* can recombine independently of FtsK, hinting at a role in plasmid dimer resolution.

### *xrs*-cassettes found on *A. baumannii* plasmids do not form excisable modules

Because *xrs* are frequently found flanking adaptative genes and because related *Ab* plasmids exhibit variability for sequences flanked *xrs*, it was proposed that pairs of *xrs* form a new type of mobile element, or modules, that can be exchanged between plasmids (17, 18, 21). These modules could excise from a plasmid and integrate into another one using XerCD_Ab_ recombination. To test this hypothesis, we assayed module excision in an *E. coli* strain deleted for *xerC* and *xerD* and expressing different version of the *xerC* and *xerD* genes from a plasmid (Fig. 4A). These strains were transformed with plasmids (S) in which the *lacI* gene is flanked by the assayed pair of *xrs* in direct orientation. A deletion product (P) was observed between repeated *dif_Ab_* sites in strains expressing *xerCD* genes, either *E. coli* or *A. baumannii* ones (Fig. 4B). This shows that *dif_Ab_* can be recombined by XerCD either from *E. coli* or from *Ab* and that *E. coli* FtsK can activate XerD_Ab_ (lanes CD_Ab_ and CD_γAB_). Recombination was not detected when the strain expresses a mutated (loss of catalytic activity) version of *xerD_Ab_* (Fig. 4B, lane CD*_Ab_).

**Figure 4:**
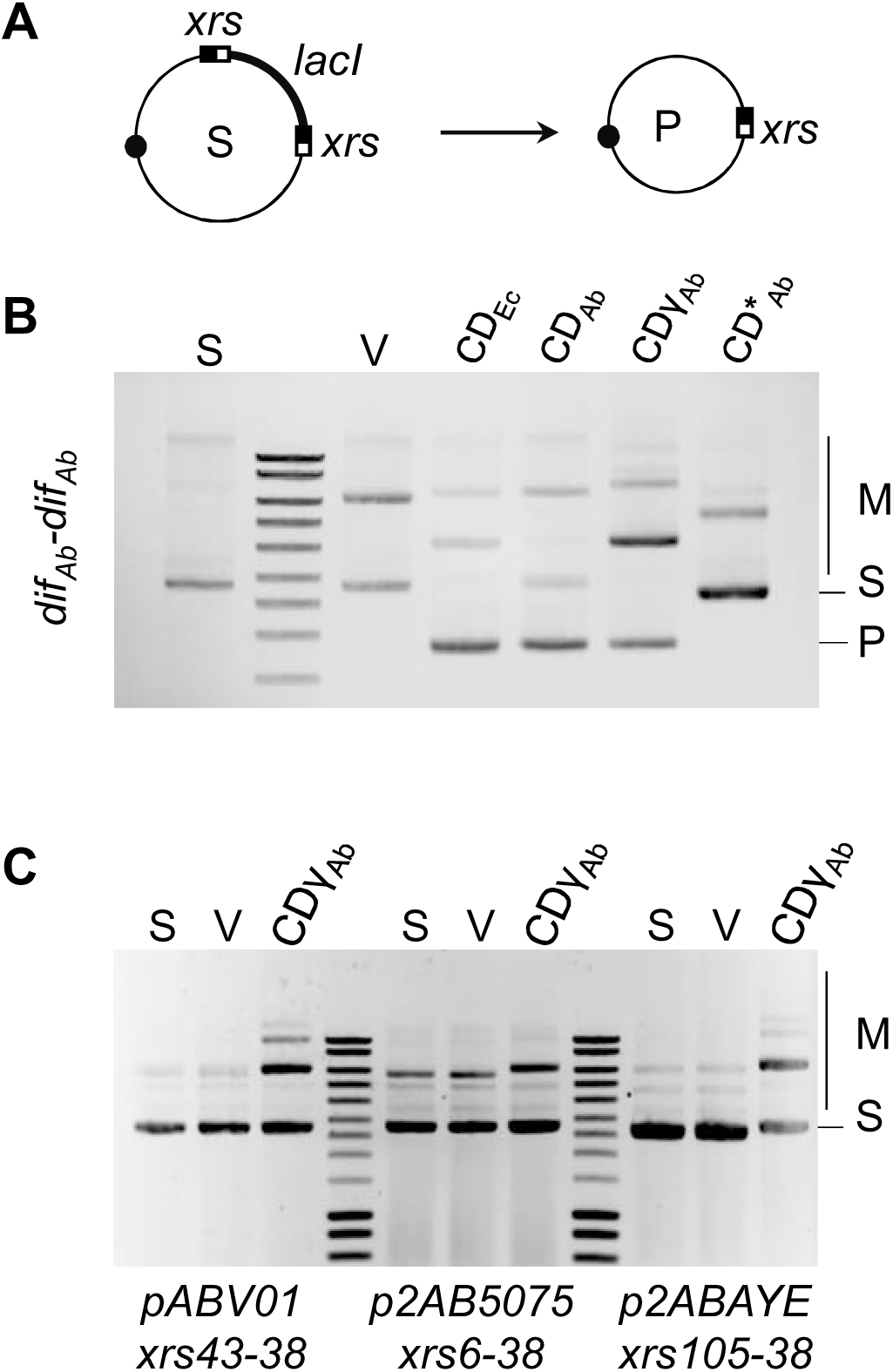
*xrs*-cassette deletion. **A)** The XerCD recombinase, when produced, should delete the *lacI* gene from the substrate plasmid (S) to give a deleted plasmid (P) and a non-replicative circle containing *lacI*, lost during divisions. **B)** Gel electrophoresis of the plasmid extraction after overnight culture of different strains transformed with a substrate plasmid containing a *dif_Ab_-lacI-dif_Ab_* cassette: S, substrate plasmid used for cells transformation; V, plasmids extracted from a strain which does not express any *xer* genes; CD*_Ec_*, plasmids extracted from a strain expressing *xerCD* from *E. coli ;* CD*_Ab_*, plasmids extracted from a strain expressing *xerCD* from *Ab;* CDg*_Ab_*, plasmids extracted from a strain expressing *xerC_Ab_* and a fusion between *xerD_Ab_* and g domain *of ftsK_Ab_.* Substrate (S) and product (P) plasmids are indicated, as well as multimeric forms of substrate and product plasmids (M). **C)** Gel electrophoresis of the plasmid extraction after overnight culture of different strains transformed by substrate plasmids (S) containing different *xrs-lacI-xrs* cassettes. The *xrs43-38* cassette represents the cassette found in pABV01, the *xrs06-38* cassette represents the cassette found in p2AB5075, and the *xrs105-38* cassette represents the cassette found in p2ABAYE,

We then assayed pairs of *xrs* mimicking genetic organization found in the pABVO1, p2AB5075 and p2ABAYE plasmids (Fig. 4C). No deletion was detected using any combination of module and recombinases. This strongly suggest that the *xrs* found in *Ab* plasmid do not form excisable modules.

Interestingly this experiment also revealed that the expression of *xerCDγ_Ab_* in these strains give rise to the formation of multimeric form of the *xrs*-containing plasmids (Fig. 4C). This confirms that *xrs* are recognized end recombined in vivo by XerCDγ*_Ab_*, leading to recombination between plasmid molecules (intermolecular), rather than within the plasmids (intramolecular).

### High recombination rate between *xrs*-carrying replicons in *A. baumannii*

Xer recombination usually allows excision of a DNA fragment flanked by pairs of *xrs* in direct orientation with identical central region (2, 15). Noteworthy, in *Ab* plasmids most pairs of *xrs* flanking adaptative genes are frequently in opposite orientation and/or have diverging central regions (18, 19, 21). This latter observation rules out intramolecular Xer recombination of Ab plasmid carrying *xrs*. However, gene exchange may results from recombination between *xrs* carried by co-existing plasmids (20). Recombination between compatible *xrs* with closely related central regions would co-integrates plasmids, allowing the exchange of genes by resolution of the co-integrate at a second pair of compatible *xrs*.

To assay intermolecular recombination between different plasmids, we took advantage of the ABAYE strain containing two different Rep_3-family plasmids that both carry multiple *xrs*: p1ABAYE (p1) and p2ABAYE (p2) (Fig. 5). Central regions of *xrs6* (p1) and *xrs105* (p2) are identical, as are the ones of *xrs107* (p1) and *xrs10* (p2), whereas the central regions of *xrs66* and *xrs38* diverge at one position. We monitored intermolecular recombination events using a PCR based approach, using pairs of oligonucleotides amplifying the DNA region resulting from intermolecular recombination between *xrs*. Recombination was observed between two sets of *xrs* (*xrs6/105* and *xrs66/38*, Fig. 5: lanes 2 & 4) but not for *xrs107/10* (Fig. 5: lane 6). Sequencing data demonstrated that recombination occurred within the *xrs* (Fig. Sup2) whereas recombination was abrogtated in a Δ*xerC* derivative (Fig. 5: lanes 7-12). Note that *xrs107* was poorly recognized by XerCD*Ab* (Fig. 2), which may explain the absence of recombination. Site specific recombination was not observed between *xrs* that does not share closely related central region (Fig. Sup2). Therefore, *Ab* plasmids form co-integrates structures by recombination between *xrs* provided closely related central regions.

**Figure 5:**
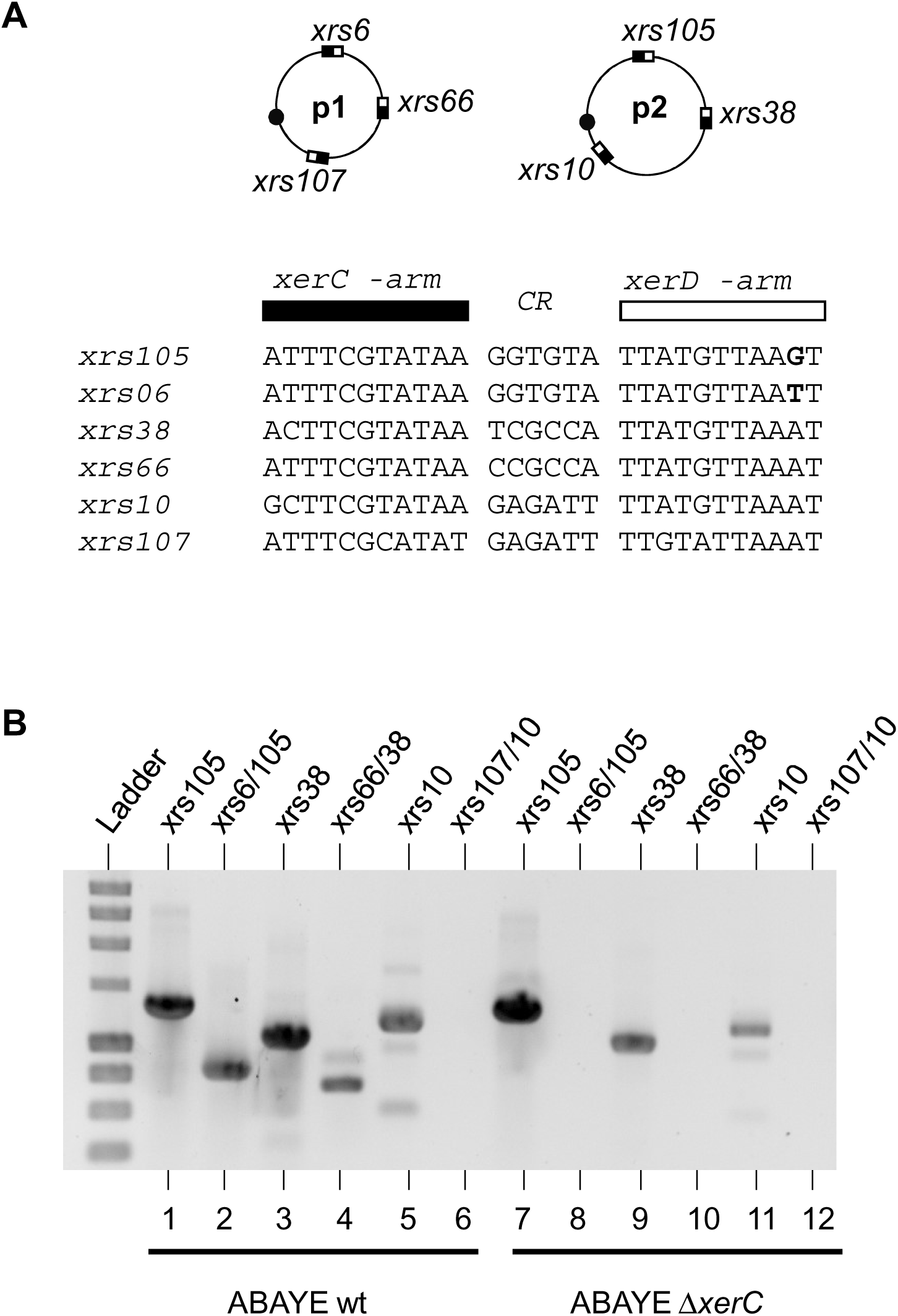
Xer-dependent plasmid recombination. **A)** simplified genetic map of p1ABAYE (p1) and p2ABAYE (p2). The *xrs* present on these plasmids are indicated and their sequences are aligned. Small black square on plasmid represents the Rep_3 replication origin of these plasmids **B)** Gel electrophoresis of the PCR reactions performed on plasmid DNA extracted from wt or D*xerC* ABAYE strains. Lanes correspond to the amplified *xrs*. For instance: lane *xrs105* corresponds to the PCR product obtained when the two primers flank *xrs105* (p2); lane *xrs6/105* corresponds to the PCR product obtained when one primer flanks *xrs6* (p1) whereas the other flanks *xrs105* (p2).

We attempted to quantify the ratio of co-integrates using ddPCR (Fig. SUP3). Co-integrate formed by *xrs6* - *xrs105* recombination represented 0.25% of the DNA matrix leading to the PCR observed from p2 (Fig. 5B lane 1). Considering that p2 appears ≈ 1,5 time more abundant than p1, we estimate a rate of co-integrate of 0.45% for p1. Consistent results were obtained for recombination between *xrs38* and *xrs66*, confirming a steady-state rate of .1 to 1% of co-integrate between the two plasmids.

## Discussion

When *xrs*-cassettes were first described in *Ab* plasmids (17), authors immediately proposed that the Xer system may be involved in the mobility of these genetic structures. However, given that *xrs* flancking each cassette have different central regions and that they are organized as inverted sequences, it was not clear how the Xer system may ensure the mobility of these *xrs*-cassettes.

We first characterized the Xer system of *Ab* to describe a highly canonical chromosome dimer resolution system (2). It catalyzes site specific recombination between *xrs* sharing the same central region, in a FtsK dependent manner. This is yet another example of the remarkable conservation and control of the Xer system among bacteria (31, 34).

Furthermore, while deletion is indeed observed between directly repeated *dif_Ab_* sequences, this phenomenon does not occur between *xrs* arranged as they are on *Ab* plasmids. This invalidates the hypothesis that the Xer system of *Ab* (*i.e.:* XerC_Ab_, XerD_Ab_ and FtsK_Ab_) would catalyze the mobility *xrs*-cassettes from one plasmid to another by a simple excision/insertion mechanism.

Our results rather suggest that *xrs* present on different *Ab* plasmids recombines between each other, leading to the formation of cointegrates. Such cointegrate of Rep_3 plasmids in *Ab* were described in previous work when plasmid encounter is triggered by their artificial transfer (19, 22). In this work, we have used the ABAYE strain which contains two Rep_3 plasmids (p1ABAYE and p2ABAYE), each containing multiple *xrs*. All the tested *xrs* are active, except *xrs*107 (p1ABAYE). This *xrs* is however mutated in the highly conserved motif “ATAA-Central Region-TTAT”, which seems to be essential for Xer binding and cutting activity (5). Two pairs of *xrs* appeared to be compatible for recombination: *xrs6* (p1ABAYE) and *xrs105* (p1ABAYE), which share de GGTGTA central region; *xrs66* (p1ABAYE) and *xrs38* (p1ABAYE), which share de TGGCGA/G central region. This suggests that *xrs* are compatible when their central region regions are identical or closely related, consistent with the known Xer recombination mechanism (2, 35). This shall be taken into account for future *xrs* annotation on *Ab* plasmids in order to predict all the possible recombination events between *Ab* plasmids.

We measured less that 1% recombination rate between active and compatible *xrs* on p1- and p2ABAYE. This could be considered low compared to Xer activity in the context of plasmid dimer resolution system where 100% of dimers are resolved (15). However, if plasmid dimer resolution described previously depends on XerCD, it is not activated by FtsK. Plasmid dimer resolution is activated thanks to the binding of DNA proteins (i.e.: PepA, ArgR or ArcA) upstream of the *xrs* allowing it to be independent of the bacterial cell cycle (2, 15). Here, we show that recombination between *xrs* found on *Ab* plasmids depends on XerCD_Ab_ but also on FtsK_Ab_ activation. As FtsK_Ab_ is only present at the septum when cells divide, recombination between xrs would only occur in the vicinity of the septum of dividing cells, explaining why we only observe few cointegrates (36, 37).

Taken together, our results show that co-integrates are dynamics entities occurring between co-residing plasmids in *Ab*, in a XerCD system-dependent manner. This strongly support a mechanistic model where Xer recombination between one pair of active and compatible *xrs* present on two different plasmids lead to cointegrate formation and that resolution of this cointegrate by recombination between another pair of active and compatible *xrs* lead cassette exchange between these plasmids (20).

Although we observed recombination between each pair of active and compatible *xrs*, we did not observe any *xrs*-cassette exchange between these plasmids. This suggest that cointegrates are more frequently formed and resolved by recombination between the same pair of *xrs* than between different pairs of *xrs*. The *xrs*-cassette exchange may thus be too rare to be detected in our experiments (30 generations per experiment), but sufficiently frequent to explain the variability observed in *Ab* plasmids in the different *Ab* strains, exposed to different selective pressure. Furthermore, p1- and p2ABAYE coexist in the same strain (ABAYE) and may have reached a stable state. It is not clear if a cassette swap between these plasmids has any advantage for the ABAYE cell. In the future, experiment will have to be designed to address this question. For instance, it could be interesting to mutate maintenance genes of p1- or p2ABAYE to destabilize them and be able “select” for an eventual *xrs*-cassette exchange.

This work demonstrates that *Ab* Rep_3 plasmids harbor multiple active Xer recombination sites and raises questions about stability of these plasmids. Indeed, if intermolecular recombination between compatible *xrs* present on different plasmid molecules leads to cointegrate formation (this work, (19, 22)), it will also lead to multimerization of these plasmids, which may be deleterious for their stability (14). This multimerization of Rep_3 plasmids have not been reported and we have not detected it in our studies. This suggests that even though compatible *xrs* present on different plasmids results in cointegrates, recombination between compatible *xrs* present on different copies of the same plasmid is somehow inhibited. It could be possible that *Ab* Rep_3 plasmids carry a dimer resolution system, like described for pSC101 or pColE1 in *E. coli* (15, 38). In the plasmid investigated in this study, this later mechanism could potentially explain the activity of *xrs38* and *xrs66*. Both of these *xrs* appear to be recognized and cleaved by both XerC_Ab_ and XerD_Ab_, even in the absence of FtsK, aligning with one of the characteristics of plasmid dimer resolution sites (15). This hypothesis will have to be tested in the future.

Taken together, these results bring experimental evidence of gene exchange between Rep_3 *Ab* plasmids could occur via a unique cointegration-resolution mechanism. It would be interesting to know if this mechanism has only been evolved by Acinetobacter and what are the advantage of such a system, for the plasmids and/or for the host bacteria.

## Acknowledgments

We thank all the members of our teams, F-X Barre, S. Venner, T. Koffel, M. Bruto and E. Accacia for helpful discussions, S. Nejmi and S. Quertinmont for technical assistance. This work was funded by the CNRS, University of Toulouse 3 – Paul Sabatier and ANR-AAPG2021-InXS. CB was supported by a fellowship from the Université de Lyon.

## AUTHOR CONTRIBUTIONS

Conceptualization: CB, XC, FC and PR. Methodology: all authors. Writing: CB, XC, FC and PR. Funding Acquisition: XC, FC and PR. Blanchais et al., Figure 1

---

**Table Sup2:** Deletion rate of a *dif_Ab_-lacI-dif_Ab_* cassette inserted in the *E. coli* chromosome at *dif* (Deghorain et al. 2011). A) The strain is deleted for *xerC* and complemented with pLN1::*xerCD_Ab_.* B) The strain is deleted for *xerC* and complemented with pLN1::xerCD*γ_Ab_*.

| Assay | Colonies |  |  | Recombination rate % |
| --- | --- | --- | --- | --- |
|  | Blue | Total | % |  |
| 1 | 46 | 440 | 10 | 10 |
| 2 | 13 | 136 | 10 |  |
| 3 | 246 | 2387 | 10 |  |

**B**
| Assay | Colonies |  |  | Recombination rate % |
| --- | --- | --- | --- | --- |
|  | Blue | Total | % |  |
| 1 | 46 | 87 | 53 | 47 |
| 2 | 209 | 498 | 41 |  |
| 3 | 232 | 507 | 46 |  |

**Figure Sup 1:**
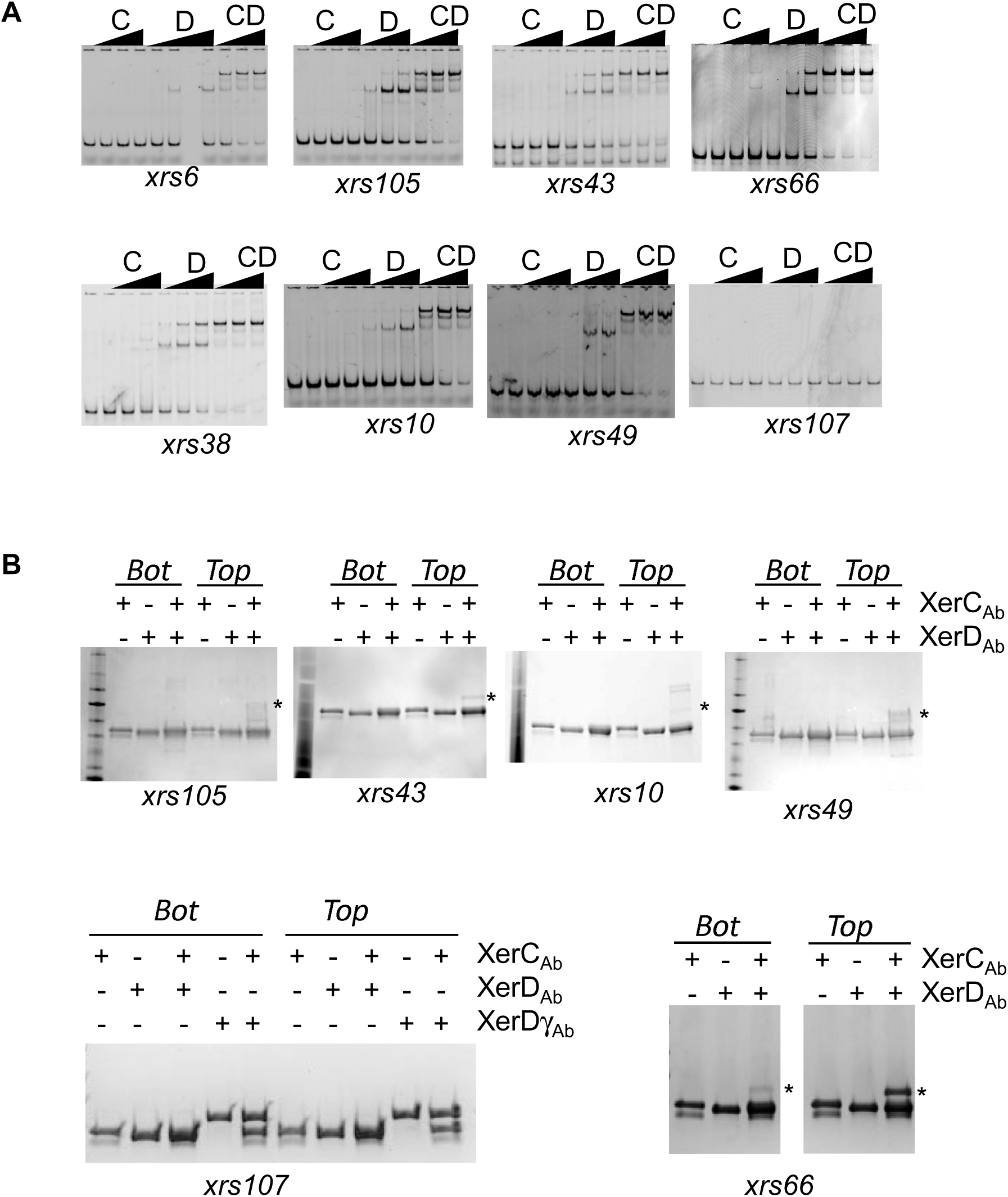
Activity of the *xrs* of *Ab (summarized in* Figure 3. **A)** EMSA of *xrs* by XerCDAb. The DNA fragment (*xrs*) is 56 bp long and is 5’-CY3. XerCAb (μM) = 0; 0.48; 1.2 and 2.4. XerDAb (μM) = 0; 0.64; 1.6 and 3.2. **B)** SDS-PAGE analysis of cleavage assays (top diagram). DNA molecules (*xrs* Top or Bottom nicked) are 10μM whereas proteins are 2 μM. Covalent products made between DNA and proteins are indicatited (*).

**Figure Sup2:**
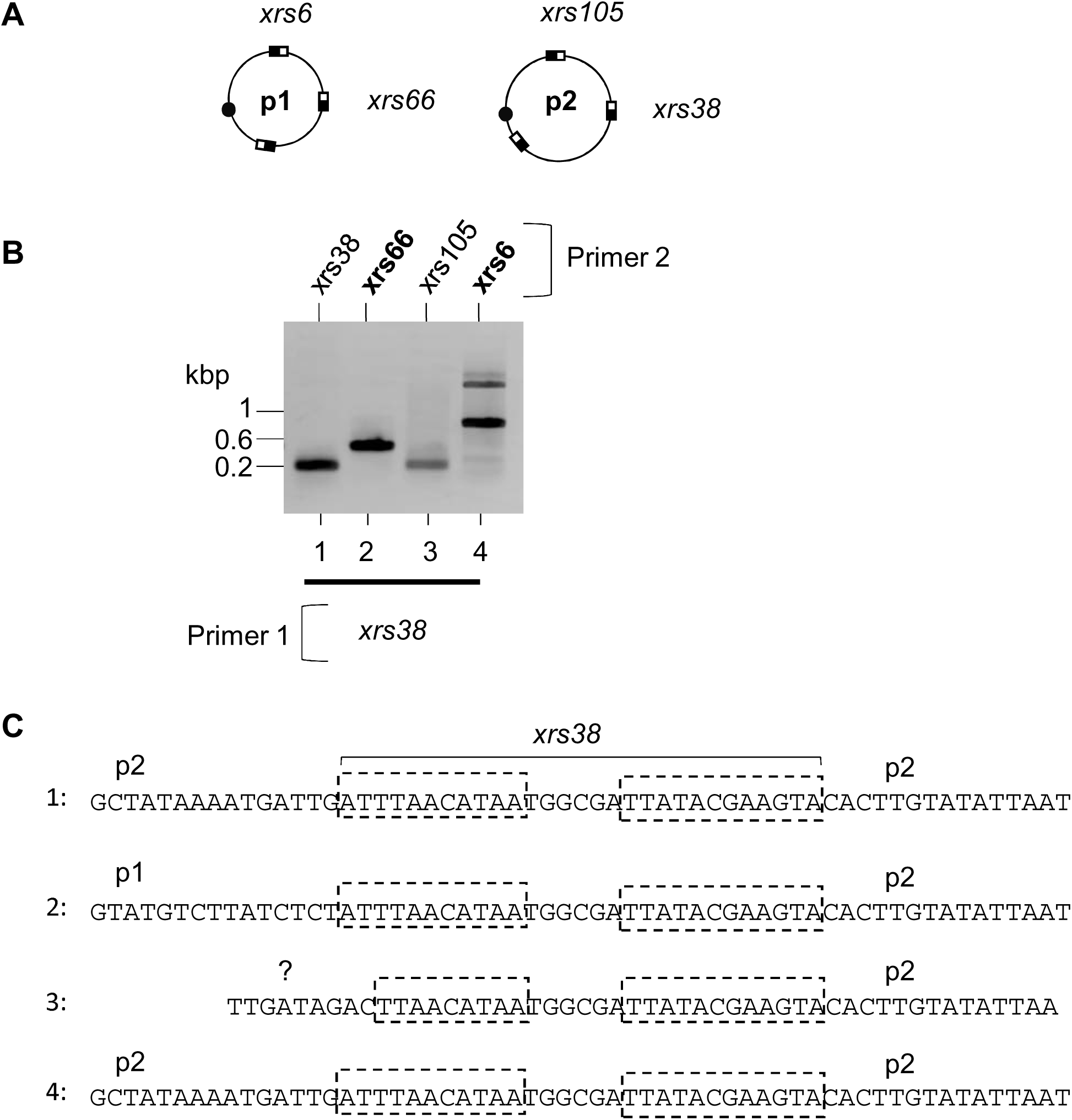
Xer-dependent plasmid recombination implicating *xrs38* (p2ABAYE). **A)** Simplified genetic map of p1ABAYE (p1) and p2ABAYE (p2). **B)** Gel electrophoresis analysis of the PCR reaction performed on plasmid DNA extracted from ABAYE strains. First primer hybridizes upstream of the *xerC-arm* of *xrs38*. The second primer hybridizes upstream of the *xerD-arm* of *xrs105* or *xrs6* or *xrs38* or *xrs66*. **C)** Sequencing revealed that PCR products of lane 1 corresponds to a non recombined p2ABAYE and that PCR products of lane 2 corresponds to recombination between p1 and p2ABAYE within *xrs38.* PCR product obtained lane 3 and 4 does not correspond to recombination between *xrs38* and one of the other tested *xrs*.

**Figure Sup 3:**
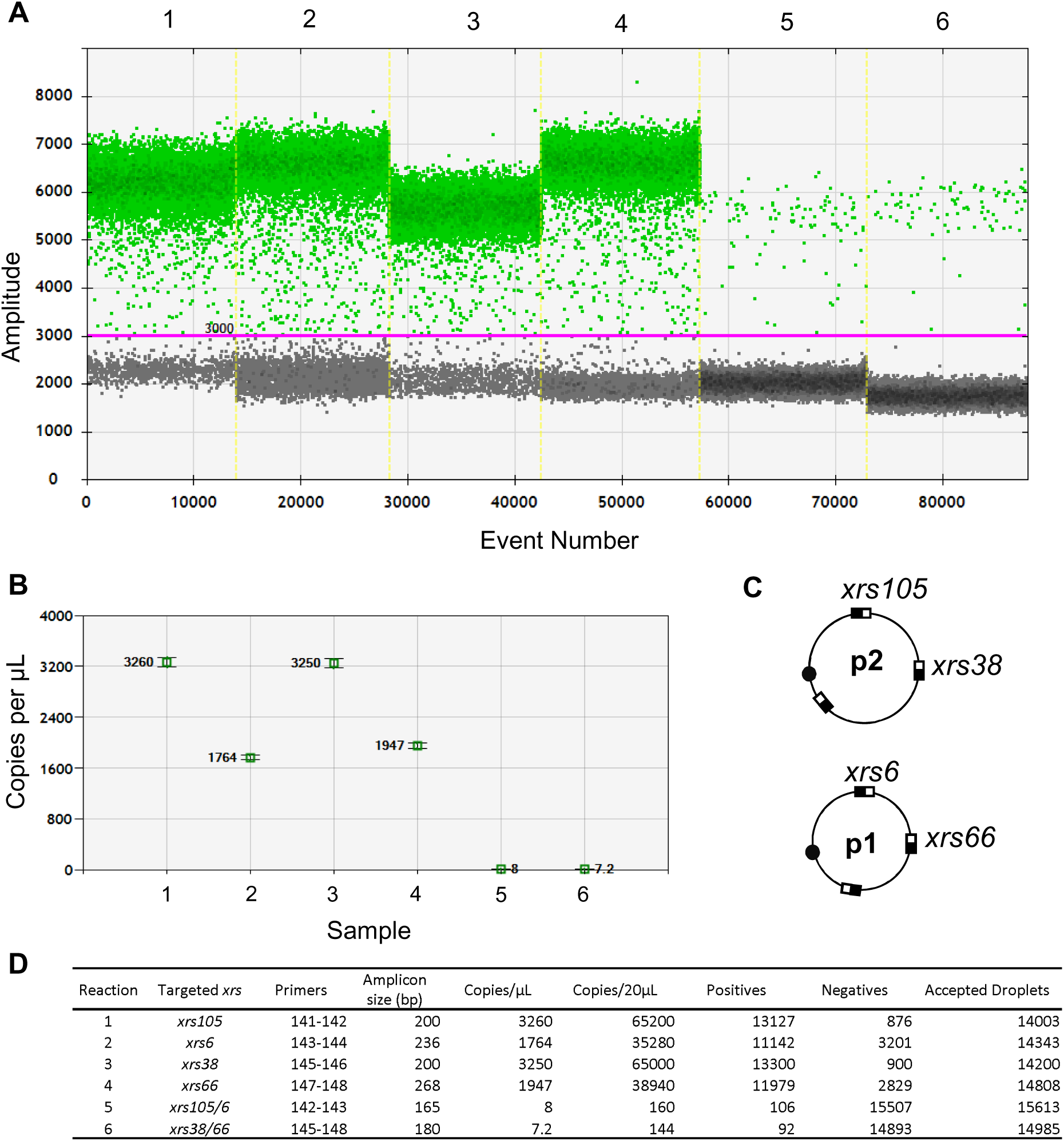
ddPCR for recombination quantification. **A)** Each columns represents single ddPCR amplification. Green dots represent positive droplets whereas black dots represent negative droplets. Column 1 corresponds to the amplification of *xrs105,* column 2 corresponds to the amplification of *xrs6,* column 3 corresponds to the amplification of *xrs38,* column 4 corresponds to the amplification of *xrs66,* column 5 corresponds to the amplification of a fusion between *xrs105* and *xrs6* (recombination between p1 and p2), column 6 corresponds to the amplification of a fusion between *xrs38* and *xrs66* (recombination between p1 and p2). **B)** The estimated copy number of each DNA matrix molecule is represented. **C) S**implified genetic map of p1ABAYE (p1) and p2ABAYE (p2). **D)** Summary of the data collected thanks to these 6 different ddPCR reactions. The estimated copy number of the matrix molecules (either recombine or not) is calculated using the Biorad QuantaSoft droplet reader software. The recombination ratio (see text) is calculated by dividing the number of recombined molecules by the sum of recombined and non-recombined molecules.

